# Rhizosphere microbes influence host circadian clock function

**DOI:** 10.1101/444539

**Authors:** Charley J. Hubbard, Robby McMinn, Cynthia Weinig

**Affiliations:** Department of Botany, University of Wyoming, Laramie, WY, USA; Program in Ecology, University of Wyoming, Laramie, WY, USA; Department of Molecular Biology, University of Wyoming, Laramie, WY, USA

## Abstract

The circadian clock is an important determinant of individual fitness that is entrained by local conditions. In addition to known abiotic inputs that entrain the circadian clock, individual pathogenic soil bacteria affect the circadian period of plant hosts. Yet, in nature, plants interact with diverse microbial communities including hundreds to thousands of microbial taxa, and the effect of these communities on clock function remains unclear. In *Arabidopsis thaliana*, we used diverse rhizosphere inoculates and both wild-type and clock mutant genotypes to test the effect of complex rhizosphere microbial communities on the host circadian clock. Host plants with an intact rhizosphere microbiome expressed a circadian period that was closer to 24 hrs in duration and significantly shorter (by 60 minutes on average) relative to plants grown with a disrupted microbiome. Wild-type host genotypes differed significantly in clock sensitivity to microbiome treatments, where the effect was most pronounced in the Landsberg *erecta* genotype and least in the Columbia genotype. Rhizosphere microbes collected from a host genotype with a short-period phenotype (*toc1-21*) and used as inoculate significantly shortened the long-period phenotype of the *ztl-1* clock mutant genotype. The results indicate that complex rhizosphere microbial communities significantly affect host clock function.

## Introduction

Across all domains of life, organisms have evolved endogenous time-keeping mechanisms known as circadian clocks [1]. In plants, the clock regulates a large percentage of the transcriptome, the metabolome, leaf-level and organismal physiology, and even the composition of beneficial rhizosphere microbial communities [2–5]. There are several well-described abiotic and biotic factors leading to clock entrainment, including photoperiod, light intensity, temperature, and herbivory [3, 6]. A recent study has shown that soil microorganisms may be another environmental factor that entrain the plant circadian clock [7]. Specifically, Zhang *et al*. (2013) found that *Arabidopsis thaliana* plants infected with *Pseudomonas syringe* had significantly shorter periods, that is, the duration of one circadian cycle, than uninfected plants. Zhang et al. (2013) demonstrate that individual microbial taxa affect clock function. In nature, plants interact with complex microbial communities consisting of hundreds to thousands of taxa [8]. Thus, the question remains as to how natural, complex microbial communities may influence clock function.

Microbiomes affect many aspects of host performance [8]. Although an organism’s own genotype has been considered as the primary determinant of phenotype for many traits, there is growing evidence that host-associated microbial communities and their genomic functions also contribute to host phenotypes. In some cases, the phenotypic effects of microbial associates exceed the effects of host genetics [8, 9]. For instance, studies of human-gut microbiome interactions have shown that the composition of gut microbial communities is a better predictor of host obesity than is host genetics [10]. The genetic pathways underlying clock function are well-described in plants, yet the magnitude of microbial effects *vs.* plant genotype on clock phenotypes is unknown. Comparing the effects of diverse inoculation treatments and host genotypes would reveal relative microbial *vs*. host genetic contributions to clock phenotypes.

In the current study, we test the effects of complex rhizosphere microbial communities on host clock function. We first characterize the effects of rhizosphere microbes on the host plant clock by comparing the circadian period of plants grown in intact *vs*. disrupted rhizosphere microbiome treatments. Next, we examine the relative effects of distinct microbial treatments *vs*. plant genotypes on circadian period. We then test if rhizosphere microbial communities found in association with host plants with short (20-hr) *vs*. wild-type (24-hr) circadian periods can rescue (shorten) the long-period phenotype (28-hr) of a mutant genotype with disrupted clock function.

## Materials and Methods

### Plant material and growth conditions

To investigate the influence of rhizosphere microbes on host plant circadian period, we used *Arabidopsis thaliana* accessions that harbor the reporter gene *LUCIFERASE (LUC)*, the gene responsible for bioluminescence in the presence of the substrate luciferin in fireflies (*Photinus phralis*), linked to the native circadian gene, *COLD CIRCADIAN RHYTHM RNA BINDING 2 (CCR2)*, allowing for quantification of circadian parameters [11]. Genotypes included in the current study are Columbia (Col), Strand (Nrw), Wassilewskija (Ws), Landsberg *erecta* (L*er*), and both the short-period mutant *TIMING OF CAB EXPRESSION 1* (*toc1-21*) and the long-period mutant of *ZEITLUPE* (*ztl-1*) in the Col background.

Seeds were surfaced sterilized using a 10% bleach and 0.1% Tween solution, planted into 96- well plates containing Murashige and Skoog mineral plant growth media containing 30g/L sucrose, and cold stratified for 7 days at 4°C. To germinate and entrain plants, plates of planted seeds were placed in a Percival PGC-9/2 growth chamber (Percival Scientific, Perry, IA, USA) set on a 12 / 12 light / dark cycle at 22°C. After 7 days, plants were inoculated with 10μl of a microbial inoculant created combining 10g of soil with 90ml of ROH_2_O and filtered through 1 000, 500, and 212 sieves to remove nematodes that could detrimentally affect plant performance [12]. 24 hours after inoculation, plates were moved into an ORCA-II ER digital camera (Hamamatsu Photonics, Bridgeport, NJ, USA) for imaging and quantification of bioluminescence.

Each 96 well plate contained 3 genotypes with 32 replicates each. Within each genotype there were 4 rhizosphere microbiome treatments with 8 replicates each. Rhizosphere microbiome treatments used in this study include: intact, disrupted, Columbia, and *toc1-21*. To generate the intact treatment, we used soil from the Catsburg site in North Carolina that has a well-documented occurrence of wild *A. thaliana* plants [13]. For the disrupted treatment, intact inoculant was autoclaved or filtered through 0.2μm mesh to disrupt microbial communities while maintaining the nutrient load found in the intact treatment. The Columbia and *toc1-21* rhizosphere microbiome treatments reflect rhizosphere microbial communities collected from the Col and *toc1-21 A. thaliana* genotypes (that had been initially treated with the Catsburg inoculant). The Col and *toc1-21* rhizosphere treatments were included to determine if microbial communities might differ in their effects on clock period, and specifically if microbial communities found in association with a short-period mutant *vs*. wild-type host might differentially affect (shorten) the circadian period of the long-period mutant, *ztl-1*, which typically expresses a circadian period of ~28hr. All experiments were replicated 2-3 times.

### Circadian Imaging

Prior to imaging plants, 20μl of 100mM D-luciferin monopotassium salt and 0.01% Triton X-100 solution was added to each well. 30-minute-long exposure images were taken every hour for 160 hours to quantify bioluminescence. Traces were then trimmed to 72hr windows and manually inspected to ensure quality [14]. Traces that failed quality control were discarded. High quality traces were then uploaded onto Biodare2 (https://biodare2.ed.ac.uk) to estimate circadian period [15]. We used Two-Way ANOVAs to characterize the effects of rhizosphere microbiome treatment, plant genotype, and their interaction on circadian period and Tukey’s Honest Significant Differences *post hoc* contrasts to characterize differences between genotypes and rhizosphere microbiome treatments.

### Replication in the wild *A. thaliana* relative *Boechera stricta*

We characterized the influence of intact *vs.* disrupted rhizosphere microbiome treatments on the period of two *B. stricta* genotypes with natural variation in clock period, genotype 15 (~22hr period) and genotype 14 (~23hr period) using the simulated fall entrainment conditions and circadian leaf movement assays described in Salmela *et al.* 2016. 15 replicates of each genotype were grown in each of two rhizosphere microbiome treatments, intact *vs*. disrupted, which were generated using soil from the genotypes’ home site (South Brush Creek, Wyoming, USA, 41.331°N, -106.504°W) and otherwise grown following the methods described above. As before, Two-Way ANOVAs were used to characterize the effects of rhizosphere microbiome treatment, plant genotype, and their interaction on circadian period.

## Results and Discussion

The rhizosphere microbiome is often referred to as an extension of the plant genome, as complex microbial communities affect many aspects of plant performance [5, 8, 9]. However, the extent to which complex rhizosphere microbial communities influence clock function remains unclear. Here, we characterized the influence of the rhizosphere microbiome on host clock function. We specifically addressed the questions, 1) do complex rhizosphere microbial communities influence host circadian period? 2) What are the relative effects of the rhizosphere microbiome *vs.* plant genotype on circadian period? And 3) can rhizosphere communities shaped by plants with short (21-hr) or wild-type (24-hr) circadian periods rescue long-period mutant phenotypes?

Recent evidence suggests that infection with a single pathogenic microbial taxon, *P. syringae* influences plant clock function through an effect on immune signaling [7]. We find that complex communities of rhizosphere microbes significantly affect host plant circadian period (**Figure 1**). Specifically, we find the circadian period is 60 minutes longer on average among plants grown with a disrupted microbiome relative to plants grown with an intact microbiome, and the effect of this clock deceleration is that period deviates increasingly from 24 hrs (**Figure 1**). Wild-type genotypes (Col, L*er*, Ws, and Nrw) differed significantly in the sensitivity of clock period to the rhizosphere microbiome, with period being unaffected (Col) or changing by as much as 96 minutes (L*er*). Likewise, in genotype 14 of *B. stricta*, we find that the period of plants grown in disrupted treatment is up to 30 minutes longer than in plants grown in the intact treatment (**Figure 2**). The largest observed changes in clock period could well affect plant fitness, because the mismatch between endogenous and environmental cycle duration can significantly affect performance [16–18] and in particular variation in clock period of this magnitude expressed in near isogenic lines affects plant performance in the field [19]. Microbial associations with eukaryotes are, of course, evolutionarily ancient, and on average appear to modulate endogenous host circadian cycles, although some host genotypes appear more sensitive than others to the presence of microbes.

**Figure 1:**
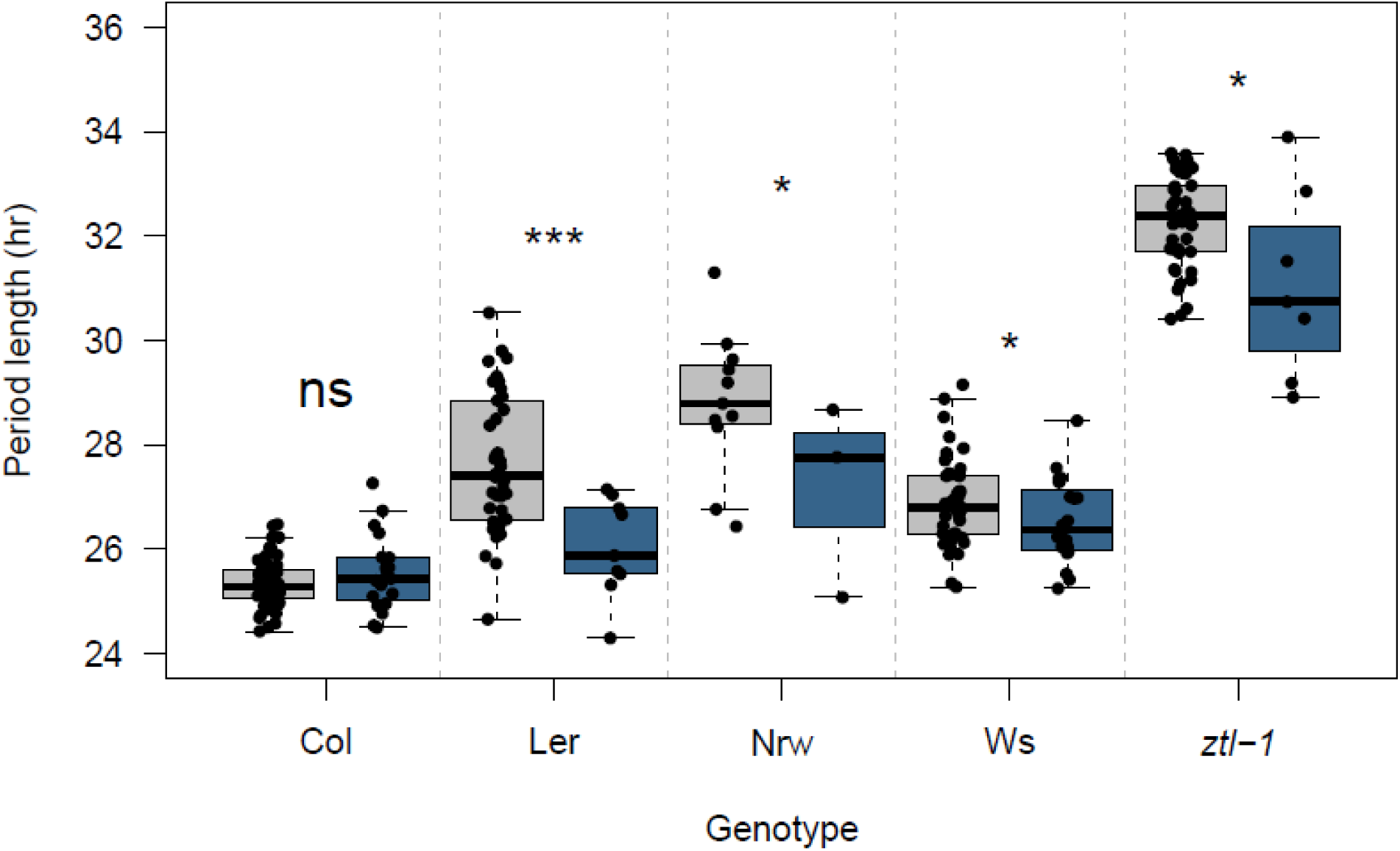
Rhizosphere microbiome treatment, intact (blue) *vs.* disrupted (gray), significantly alters plant circadian period (P < 0.001). Similarly, plant genotype (P < 0.001) and interactions between rhizosphere microbiome treatment and plant genotype (P < 0.001) significantly affect plant circadian period. The top and bottom of boxes represent the 75th and 25th percentiles, respectively. Whiskers represent 1.5 times the interquartile range. Asterisks denote significant differences between intact *vs.* two disrupted rhizosphere microbiome treatments within a given genotype: P < 0.05 *, P < 0.001 ***, ns = non-significant

**Figure 2:**
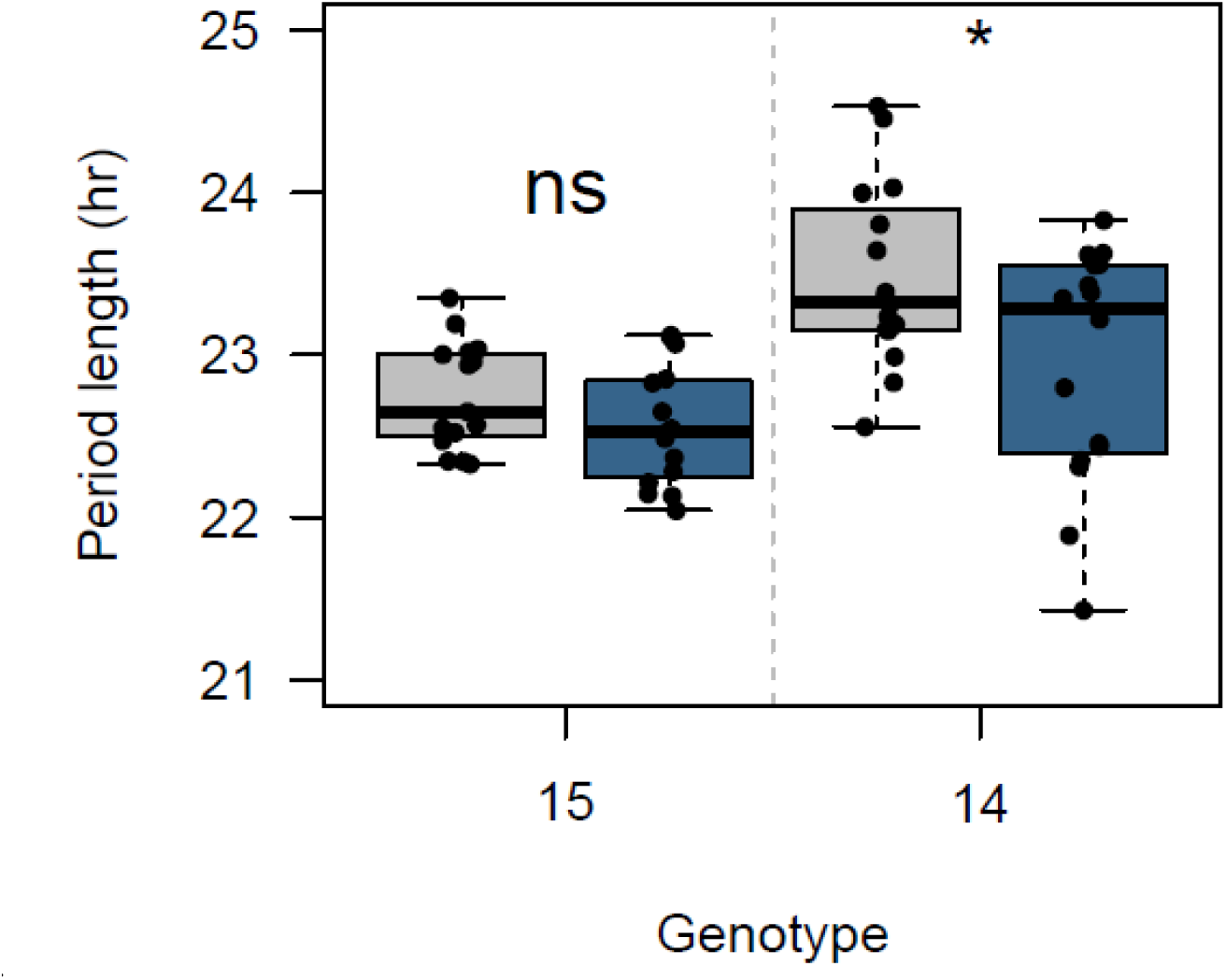
Rhizosphere microbiome treatment, intact (blue) *vs.* disrupted (gray), significantly alters the circadian period of genotype 14 (P = 0.032). Similarly, plant genotype significantly affected plant circadian period (P < 0.001). The top and bottom of boxes represent the 75th and 25th percentiles, respectively. Whiskers represent 1.5 times the interquartile range. Asterisks denote significant differences between intact *vs.* two disrupted rhizosphere microbiome treatments within each genotype: P < 0.05 *, ns = non-significant

The 1.5-hour difference in circadian period observed for at least some genotypes (L*er*) across the intact *vs.* disrupted rhizosphere microbiome treatments reflects a substantial portion of the range in circadian period expressed by the 4 wild-type *A. thaliana* accessions used in this study (50%) and by a larger panel of 150 European accessions (23%) [20]. Thus, rhizosphere microbes play an important role in clock function, but microbial effects appear of lesser magnitude than host genetic differences. As noted, there were significant interactions between rhizosphere microbiome treatment and plant genotype (P < 0.001), where the period of some genotypes (Col in *A. thaliana* and 15 in *B. stricta*) were not affected by treatment, while the period of other genotypes (L*er*, Nrw, Ws in *A. thaliana* and 14 in *B. stricta*) were more strongly responsive to rhizosphere microbiome treatments. Such genotype by rhizosphere microbe interactions suggest that there may be host genetic factors that permit microbes to affect the clock. A larger survey of diverse naturally occurring microbial communities and host genotypes will provide additional insights into the effect of microbial associates on host clock function.

Compositional differences among intact rhizosphere microbiome treatments differentially affected the circadian period of the long period clock mutant *ztl-1* (**Figure 3**). Here, *ztl-1*’s period was significantly shortened when grown in the *toc1-21* rhizosphere microbiome treatment compared to the disrupted rhizosphere microbiome treatment (P = 0.016). To estimate circadian period using bioluminescence, plants are commonly grown in sterile media amended with antibiotics like kanamycin, suggesting that period estimates are made on plants either without rhizosphere bacterial associates or with severely disrupted communities [17, 19]. As a result, such studies may not fully capture variability introduced by associated rhizosphere microbes, and the variation among that exists among plant genotypes in natural settings. Further, although rhizosphere microbiome treatments shaped by the Col and *toc1-21* genotypes do not fully rescue clock misfunction in *ztl-1*, the findings presented here suggest that rhizosphere legacy effects could affect circadian period across generations.

**Figure 3:**
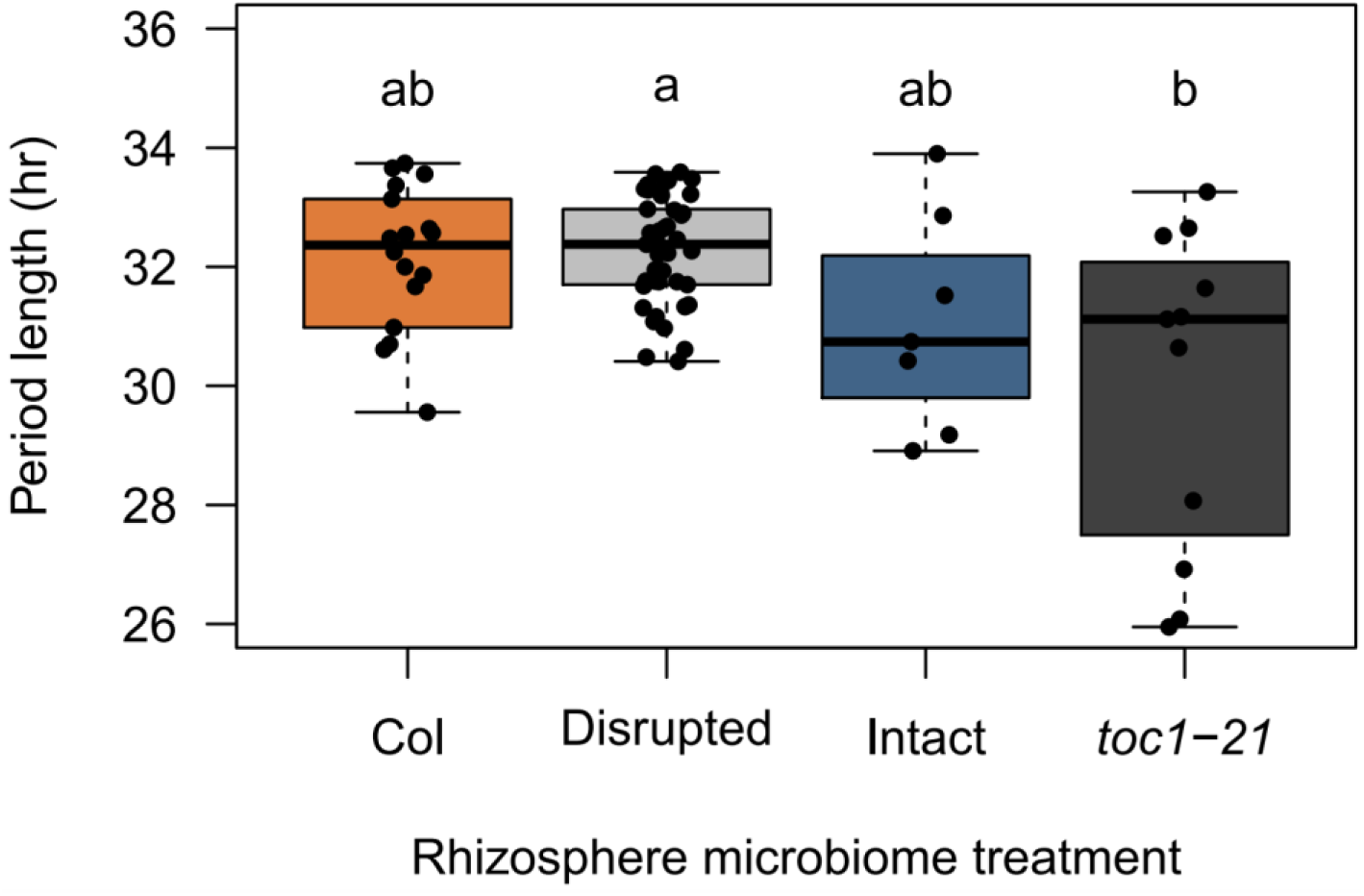
Rhizosphere microbiome treatment leads to clock deceleration in *ztl-1*. The circadian period of *ztl-1* replicates was significantly shortened when grown with rhizosphere microbial inoculate shaped by plants with short (21-hr, *toc1-21*) periods (P = 0.016) compared to *ztl-1* replicates grown with a disrupted microbiome treatment or with inoculate shaped by plants with a wild-type period (24-hr, Col). The top and bottom of boxes represent the 75th and 25th percentiles, respectively. Whiskers represent 1.5 times the interquartile range. Letters denote significant differences based on Tukey’s Honest Significant Differences *post hoc* comparisons between rhizosphere microbiome treatments.

In sum, we observed that complex microbial communities significantly influence host plant clock function. Specifically, the effect of the rhizosphere microbiome on clock entrainment and period length rivals that of differences among some plant genotypes and is equivalent to moderate changes in well-described abiotic inputs, such as a 2-hour shift in photoperiod or several degrees change in temperature [1, 3]. Further, the effect of rhizosphere microbiomes on circadian clock period is likely sufficient to alter plant performance, based on prior studies testing the effects of quantitative clock variation on plant performance [17, 19]. Given the pronounced effects intact microbes have on the plant clock, future work should attempt to identify the specific taxa and mechanisms by which microbes influence circadian period.

## Acknowledgments

This work was supported by the National Science Foundation grants IOS-1444571 and EPS-1755726 to C.W. We would like thank Matthew J. Rubin and Jenney Moua for their help on this project.

